# Balanced-detection visible-light optical coherence tomography

**DOI:** 10.1101/2021.06.08.447560

**Authors:** Ian Rubinoff, David A. Miller, Roman Kuranov, Yuanbo Wang, Raymond Fang, Nicholas J. Volpe, Hao F. Zhang

## Abstract

Increases in speed and sensitivity enabled rapid clinical adoption of optical coherence tomography (OCT) in ophthalmology. Recently visible-light OCT (vis-OCT) achieved ultrahigh axial resolution, improved tissue contrast, and new functional imaging capabilities, demonstrating the potential to improve clincal care further. However, limited speed and sensitivity caused by the high relative intensity noise (RIN) in supercontinuum lasers impeded the clinical adoption of vis-OCT. To overcome these limitations, we developed balanced-detection vis-OCT (BD-vis-OCT), which uses two calibrated spectrometers to cancel noises common to sample and reference arms, including RIN. We analyzed the RIN to achieve a robust pixel-to-pixel calibration between the two spectrometers and showed that BD-vis-OCT enhanced system sensitivity by up to 22.2 dB. We imaged healthy volunteers at an A-line rate of 125 kHz and a field-of-view as large as 10 mm × 4 mm. We found that BD-vis-OCT revealed retinal anatomical features previously obscured by the noise floor.

## Introduction

Optical coherence tomography (OCT) images the living human retina noninvasively at micrometer-scale volumetric resolutions [1, 2]. Since its first demonstration in the early 1990s, OCT has rapidly become the clinical imaging standard for diagnosing, treating, and monitoring nearly all retinal diseases [2, 3]. The clinical adoption of OCT can be partially attributed to technical advancements in imaging speed and sensitivity. Development of spectral-domain OCT enabled high-speed imaging at 100s of kHz with increased sensitivity [2–5]. Traditionally, spectral-domain OCT has operated in the near-infrared (NIR) wavelength range (800 nm-1500 nm).

The recent development of visible-light OCT (vis-OCT) [6], which operates near 500 nm – 600 nm, has shown promise for providing valuable information not available in NIR OCT. Using shorter wavelengths, vis-OCT enables an axial resolution < 2 *μ*m [6–9], 2-3-fold higher than clinical NIR systems. Combining high axial resolution with increased tissue scattering contrast from visible-light, vis-OCT reveals or enhances retinal structures previously inaccessible by NIR OCT. Recent studies have found that vis-OCT can delineate Bruch’s membrane (BM) [8–10], a structural origin of macular degeneration, and the inner plexiform layer (IPL) sublayers [11, 12], which contains scattering information from dendritic connections that may be a biomarker for glaucoma. Additionally, vis-OCT achieved retinal oximetry by analyzing spectral contrast between oxygenated and deoxygenated blood [13], opening a new window for functional retinal imaging.

Although vis-OCT enhances structural and functional information in the retina, it has unique limits that hamper its clinical adoption. Perhaps the most significant limitation is the light source. Unlike NIR-OCTs, which use low-cost, low-noise superluminescent diodes (SLDs), vis-OCT relies on supercontinuum lasers. Supercontinuum lasers have power fluctuations referred to as relative intensity noise (RIN) [6, 14, 15]. RIN imparts intensity fluctuations on each spectrometer pixel, degrading the measurement of the interference fringe. After the Fourier transform, these fluctuations are evident in increased image background noise, degrading image quality. RIN can be suppressed by increasing the exposure time of the spectrometer’s camera (decreasing A-line rate) to average out intensity fluctuations [14]. Therefore, vis-OCT researchers often limit A-line rates near or below 30 kHz in human retinal imaging [8–11, 16, 17]. Such reduced A-line rates induce two challenges. First, head and eye motions [9, 18] make optical alignment, large field-of-view (FOV) volumetric imaging, and image registrations for speckle reduction much more challenging. Second, prolonged imaging time increases light exposure in the eye. Although within ANSI laser safety standards [17], the visible-light illumination can be distracting and uncomfortable, making eye fixation challenging during alignment and acquisition. The increased eye motion and high RIN reduce vis-OCT’s capabilities in resolving minute retinal anatomical features, such as IPL sublayers and BM, and measuring functional parameters, such as oximetry. Recently, Song et al. reported vis-OCT retinal imaging at an A-line rate of 100 kHz [19]; however, they limited the illumination bandwidth to 35 nm to increase power density, which degraded axial resolution to ~ 5 μm and did not solve the RIN problem.

Until now, there has been no good solution to achieve a high A-line rate, maintain sufficient bandwidth for high axial resolution and spectroscopic contrast, and suppress RIN at the same time in vis-OCT. Our solution is to develop balanced-detection (BD) vis-OCT, which uses two calibrated spectrometers to cancel noises (including RIN) common to the sample and reference arms [3, 20, 21]. BD has been well-demonstrated in swept-source OCT [2, 3], where two single-element detectors record single-wavelength interference sweeping through the entire bandwidth to reject influences from light source energy instability and common noises. BD was also tested in spectrometer-based OCT systems [21–26]. A comprehensive study by Bradu et al. found no noise floor reduction, partially attributed to poor calibration between the two spectrometers [22]. Other demonstrations showed limited success, where researchers routinely reported 3–6 dB increases in signal intensity but no significant reduction in the noise floor [21, 23–26]. Because BD equally splits the interference signals into two separate paths before being detected by two spectrometers for combination, the reported improvement in signal amplitude is not obvious when comparing with a single spectrometer detection without splitting the interference signal [22]. One study circumvented the calibration problem by interleaving detection of fringes in time on the same spectrometer [26]. However, this degraded the effective A-line rate by 50%, limiting the key speed benefit of BD. As outlined by Bradu et al., reduction of RIN should be a key goal of BD but is particularly constrained by the inability to precisely calibrate two spectrometers.

Recently, Kho et al. [27] reported that excess noise (dominated by RIN) from a supercontinuum source is spectrally encoded on the spectrometer. This allowed for extremely accurate pixel-to-wavelength mapping of the RIN itself. We developed BD-vis-OCT and calibrated two spectrometers based on the spectrally-encoded RIN. After calibrating two spectrometers with pixel-to-pixel precision, we enhanced vis-OCT sensitivity by up to 22.2 dB and significantly reduced the noise floor. As a result, we imaged living human retinas at an A-line rate of 125 kHz for the first time and found that BD-vis-OCT revealed retinal anatomical features that were otherwise obscured by the high noise floor.

## Methods and materials

### Experimental setup

Fig. 1A illustrates the BD-vis-OCT system used for human retinal imaging. The system is based on a modified Mach-Zehnder interferometer (MZI) configuration. A supercontinuum laser (SCL, SuperK 78 MHz EXW-6, NKT Photonics, Denmark) delivered light to a multistage filter set consisting of a dichroic mirror (DM1, DMSP650, Thorlabs, NJ), a wire grid polarizer (P, WP25M-VIS, Thorlabs), a bandpass filter (BPF, FF01-560/94-25, Semrock, NY), and a spectral shaping filter (Hoya B-460, Edmund Optics, NJ). We coupled the light into a 10:90 fiber coupler (FC1, TW560R2A2, Thorlabs). The 10% output port of FS1 delivered light to the sample arm (Fig. 1A, bottom). A collimating lens (CL) collimated a 2.5-mm diameter beam incident on a galvanometer (Cambridge Technology, MA). A two-lens telescopic system (L1 and L2) with a 3:2 magnification ratio delivered 240 *μ*W to the eye. Meanwhile, a red diode laser (DL, LPS-675-FC, Thorlabs) delivered 5 *μ*W to the eye for fixation. We separated the fixation light from the vis-OCT path using a dichroic mirror (DM2, 3114-666, Alluxa, CA). A microelectromechanical scanner (MS, Mirrorcle, Richmond, CA) scanned and adjusted a star-shaped fixation pattern on the retina during vis-OCT imaging. The 90% output port of FC1 was input to a transmission-type reference arm, consisting of a polarization controller (PC), a fiber delay line (FD), and an adjustable air delay (AD) line. The AD and dispersion compensation (DC) matched the double-pass path length in the free space part of the sample arm, while the FD matched the double-pass fiber path length in the sample arm.

**Figure 1.**
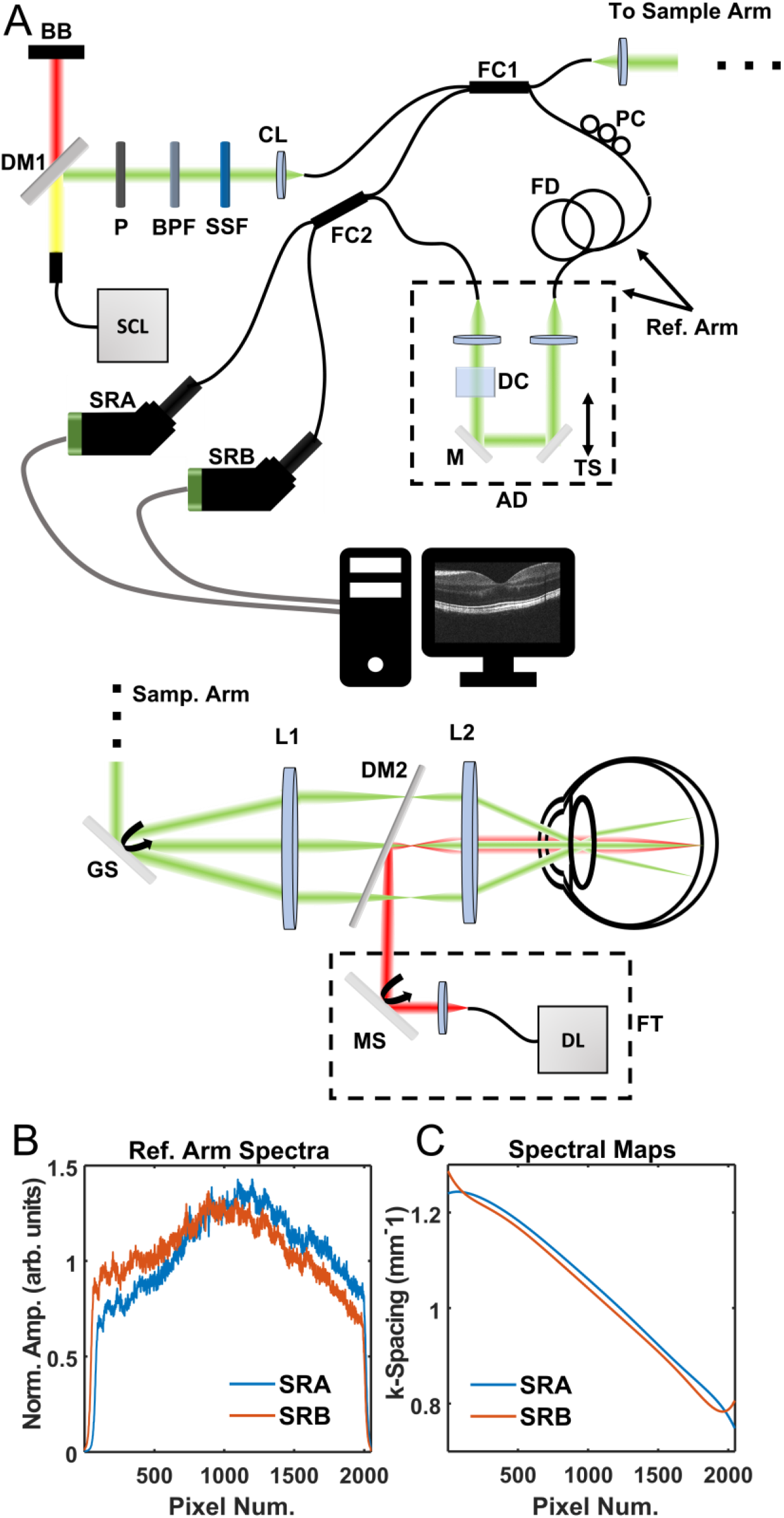
(A) Schematic of BD-vis-OCT system. SCL: supercontinuum laser, DM: dichroic mirror, BB: beam block, P: polarizer, BPF: band pass filter, SSF: spectral shaping filter, CL: collimating lens, FBS: fiber beam splitter, PC: polarization controller, FC: fiber coupler, FD: fiber delay, TS: translation stage, M: mirror, DC: dispersion compensation:, AG: air gap, SRA: spectrometer A, SRB: spectrometer B, GS: galvanometric scanner, L: lens, MS: microelectromechanical scanner, DL: diode laser, FT: fixation target; (B) reference arm spectra; (C) spectrometer spectral maps

Backscattered light from the sample arm and transmitted light from the reference arm interfered in a 50:50 fiber coupler (FC2, TW560R5A2, Thorlabs). We collected the interfered light from both output arms using two spectrometers (SRA and SRB, Blizzard SR, Opticent Inc., IL,) which offers a maximum A-line rate of 135 kHz using a high-sensitivity 1D CCD camera with adjustable gains (OctoPlus, Teledyne e2v). We subtracted the SRA detection from the SRB detection for BD. Due to manufacturing imperfection, the splitting ratio of FC2 was slightly asymmetric, as shown in Fig. 1B, which did not significantly compromise the BD performance. SRA and SRB also had slight differences in wavenumber (k) spacing with respect to the spectrometer camera pixel index, as shown in Fig. 1C. We measured an *in vivo* axial resolution of 1.7 *μ*m with SRB alone (single detection, or SD) and with SRA & SRB for BD (see **Supplementary** Fig. S1). The point-spread functions (PSFs) for SD and BD overlapped almost identically.

### Balanced detection processing

We acquired A-lines in SRA and SRB simultaneously. We loaded the respectively detected fringes and digitally matched their DC shapes to compensate for the small difference in splitting ratio of the FC2 (Fig. 1B). Then, we removed the DC components from the respective fringes. To ensure pixel-to-pixel calibration (Fig. 2), we interpolated the fringes from SRA to linear with respect to SRB using a calibration map generated by the RIN [27]. After calibration, we subtracted the respective fringes between SRB and SRA. Finally, we performed traditional image reconstruction, including k-space interpolation, compensation for dispersion mismatch, and the Fourier transform [2].

**Figure 2.**
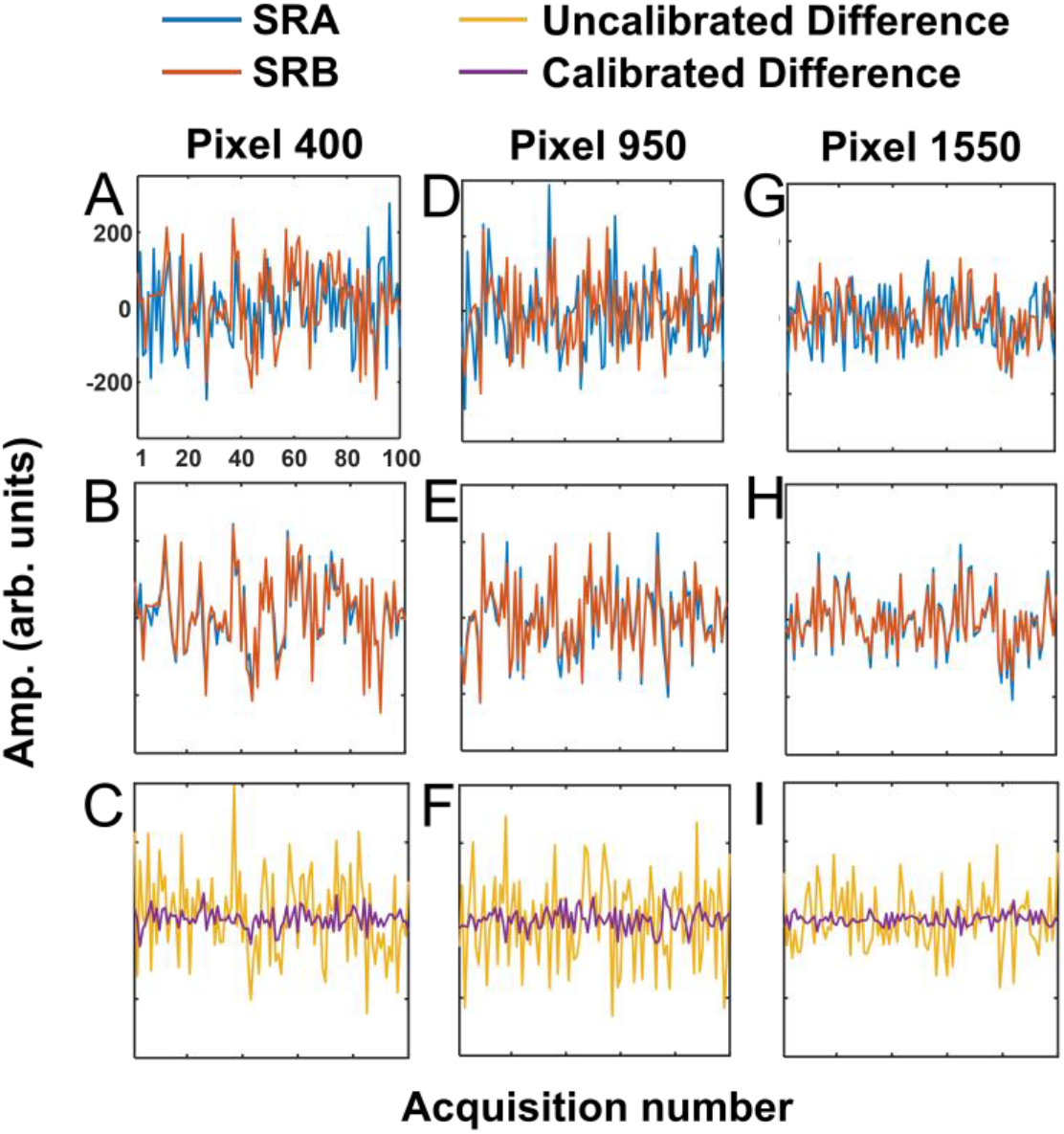
Spectrometer noises for pixel 400 without calibration (A), with calibration (B), and after subtraction between spectrometer B and A (C); (D-F) Same noises for pixel 950; (G-I) Same noises for pixel 1550.

### Phantom eyeball imaging

We imaged a phantom eyeball (OCT Model Eye, Rowe Technical Design, CA) using a rectangular scan consisting of 512 A-lines × 64 B-scans at an A-line rate of 125 kHz (7.7 *μ*s exposure time, 0.3 *μ*s readout time). We imaged at the lowest and highest camera gains.

### Human imaging

We imaged the eyes of four healthy volunteers between 25 and 47 years of age. All volunteers provided informed consent before imaging. BD imaging was performed at an A-line rate of 125 kHz (7.7 *μ*s exposure time, 0.3 *μ*s readout time) using the highest camera gain. Vis-OCT illumination light power was no higher than 240 *μ*W on the cornea. We scanned multiple patterns: small FOV with a 4 mm × 4 mm scanning range and 512 A-lines × 256 B-scans; medium FOV with a 7 mm × 4 mm scanning range and 1024 A-lines × 256 B-scans; large FOV with a 10 mm × 4 mm scanning range and 1024 A-lines × 256 B-scans; and high-density speckle reduction (HDSR) [9] with either a 12 mm × 3 mm or 8 mm × 3 mm scanning range and 32768 A-lines × 16 B-scans. HDSR scans consisted of 16 scans orthogonal to the B-scan axis [9].

### Quantifying BD improvement

As outlined in the introduction, RIN is evident in an increased noise floor and can degrade image quality. To validate that BD indeed removes RIN, we compared a reference intensity in the vis-OCT A-line to the nearby noise floor. We define this measurement here as the signal-to-noise-floor ratio (SNFR)

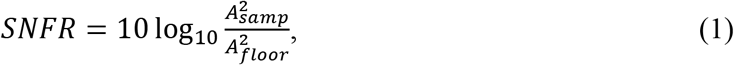

where *A*_*samp*_ is the selected amplitude of the A-line signal and *A*_*floor*_ is the amplitude of the A-line noise floor near *A*_*samp*_. Here, we compare SNFRs in single detection (SD) using only SRB and BD using both SRA and SRB. We define the SNFR gain *G* in BD as

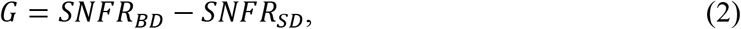

which can be rewritten as

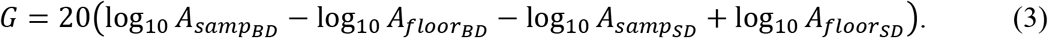

When *G* > 0, the system sensitivity increases [2]. Assuming we can achieve perfect balance and calibration between SRA and SRB, the signal gain by combining two *π*-phase shifted fringes from the two spectrometers is 6 dB [22]. The maximum *G* for BD is

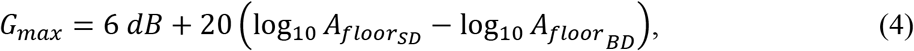

which is thusly determined by the decrease in the noise floor.

We note that for practical imaging of the human retina, 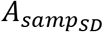 value can sometimes be very small. It has been shown that the noise floor can bias measurement of 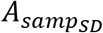 when it is sufficiently small [28]. Empirically, we found this threshold for bias was near 10 dB. To avoid this bias in the measurement of *G*, we aimed to measure *A*_*samp*_ from regions with *SNFR*_*SD*_ > 10 dB.

We also measured the contrast-to-noise-ratio (CNR), a well-accepted metric for image quality in the retina [9]. It is defined as

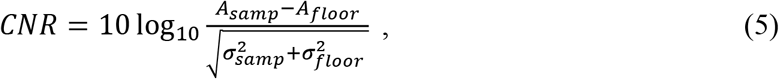

where 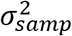 and 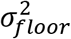 are the variances of the sample and the noise floor, respectively. For a very weak vis-OCT signal in SD, *A*_*samp*_ − *A*_*floor*_ could be very small or negative, yielding an extremely negative (< − 10 dB) or complex CNR. We considered these cases ‘noise floor limited’.

### Image metric measurements

To measure *SNFR* and *G* in the phantom eyeball, we automatically detected the brightly reflecting region at the top of the phantom (see Fig. 3). We considered the maximum amplitude at this region as *A*_*samp*_ and the average of a 20-pixel (depth) × 20-pixel (lateral) window centered 70 pixels above *A*_*samp*_ as *A*_*floor*_. We measured *SNFR* and *G* for A-lines 50 through 400 in phantom B-scans, avoiding the left and right edges, where the sample was too close to the zero-delay to accurately measure *A*_*floor*_. To measure *CNR* in the phantom retina, we used the average of a 10-pixel (depth) × 40-pixel (lateral) region below the bright line as *A*_*samp*_ and the average of a 20-pixel (depth) × 20-pixel (lateral) window centered 70 pixels above the bright line at the same lateral location as *A*_*floor*_.

**Figure 3.**
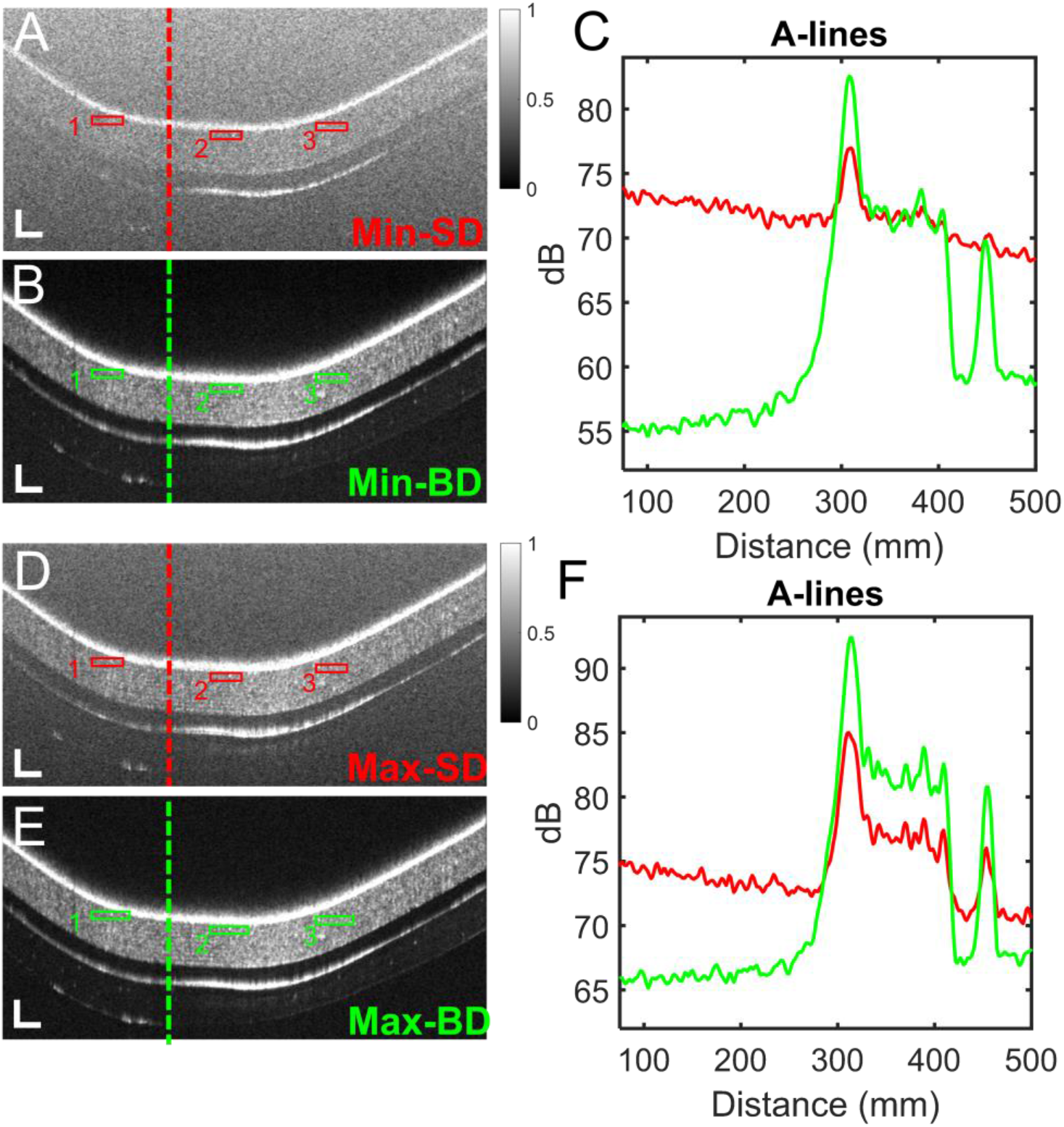
(A) Single detection and (B) balanced detection B-scans of phantom retina at lowest camera amplification level. Red and green boxes highlight measurement locations of CNR. Red and green dashed lines highlight locations of A-lines plotted in (C); (D-F) Following same layout as (A-C) but for maximum camera amplification level. B-scans magnified to show detail. Scale bars 40 *μ*m (vertical) × 225 *μ*m (horizontal) for (A, B, D, E)

To measure *SNFR* and *G* in the human retina, we identified bright reflecting regions at the internal limiting membrane (ILM) (Figs. 4–6). Since such reflections are sparse and not always visible, we consider these measurements not an objective determination of image quality but rather a validation that similar *G*s can be reached between the phantom and human retina. We considered the maximum amplitude at the ILM as *A*_*samp*_ and the average of a 20-pixel (depth) × 20-pixel (lateral) window centered 40 pixels above *A*_*samp*_ as *A*_*floor*_. We measured *SNFR* and *G* for 5 A-lines across the bright reflection. To measure *CNR* in the human retina, we used the average of a 10-pixel (depth) × 40-pixel (lateral) region in the RNFL as *A*_*samp*_ and the average of a 20-pixel (depth) × 20-pixel (lateral) window centered 50 pixels above the ILM at the location as *A*_*floor*_.

**Figure 4.**
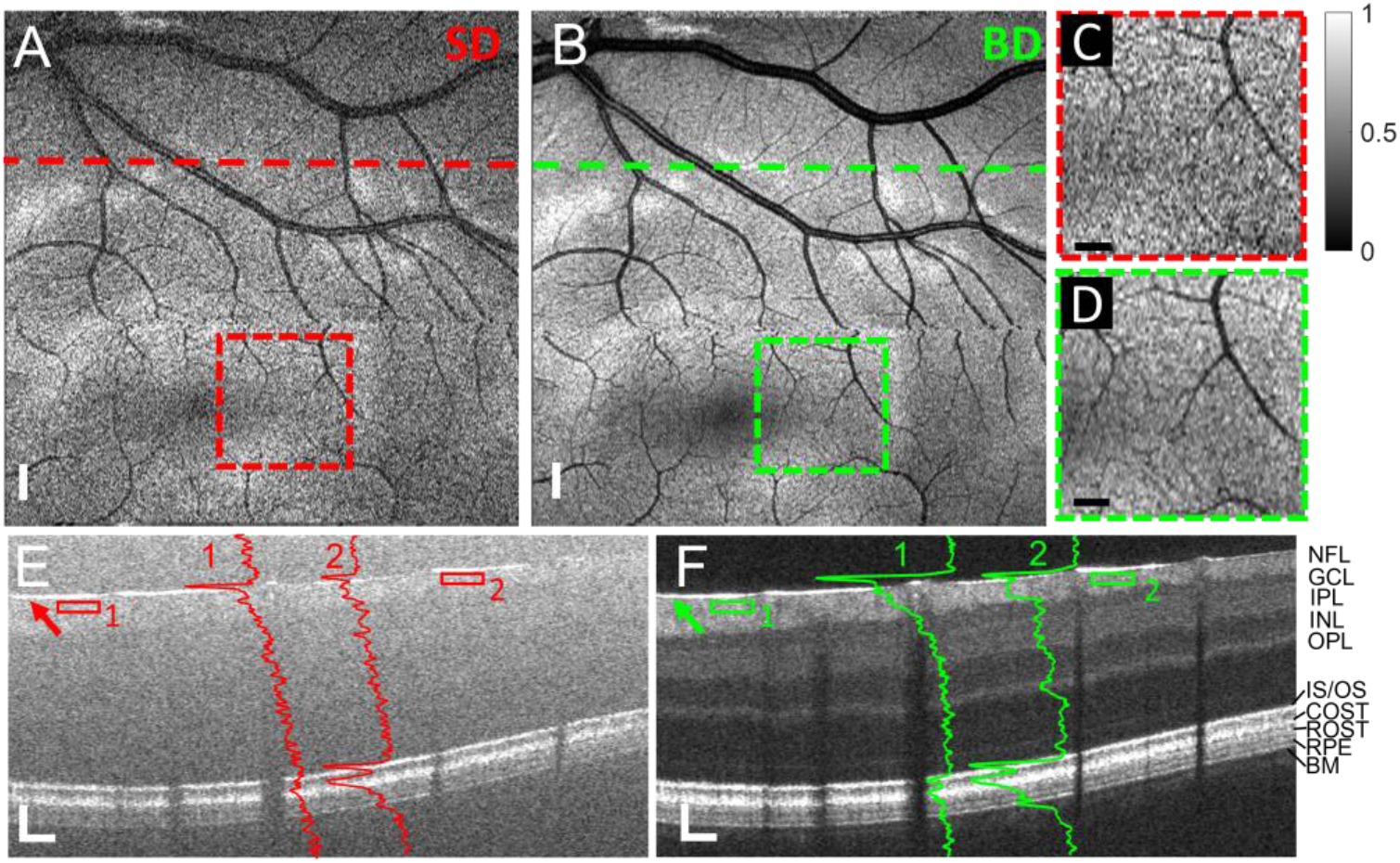
(A-F) Small field-of-view vis-OCT images of the retina of a healthy 47-year-old male (Eye 1); (A) *En face* projection near fovea for single detection; (B) *En face* projection near fovea for balanced detection; (C) Magnification from red dashed box in (A); (D) Magnification from green dashed box in (B); (E) B-scan from location of dashed line in (A); (F) B-scan from location of dashed line in (B); Red and green A-lines overlay their respective locations; Red and green arrows highlight locations of *SNFR* measurement; Solid red and green boxes highlight locations of *CNR* measurement. Scale bars in (A & B) are 275 *μ*m (isometric); scale bars in (C & D) are 150 *μ*m (isometric); scale bars in (E & F) are 60 *μ*m (vertical) × 225 *μ*m (horizontal). NFL: nerve fiber layer; GCL: ganglion cell layer; IPL: inner plexiform layer; INL: inner nuclear layer; OPL: outer plexiform layer; ELM: external limiting membrane; IS/OS: inner segment/outer segment; COST: cone outer segment tips; ROST: rod outer segment tips; RPE: retinal pigment epithelium; BM: Bruch’s membrane

**Figure 5.**
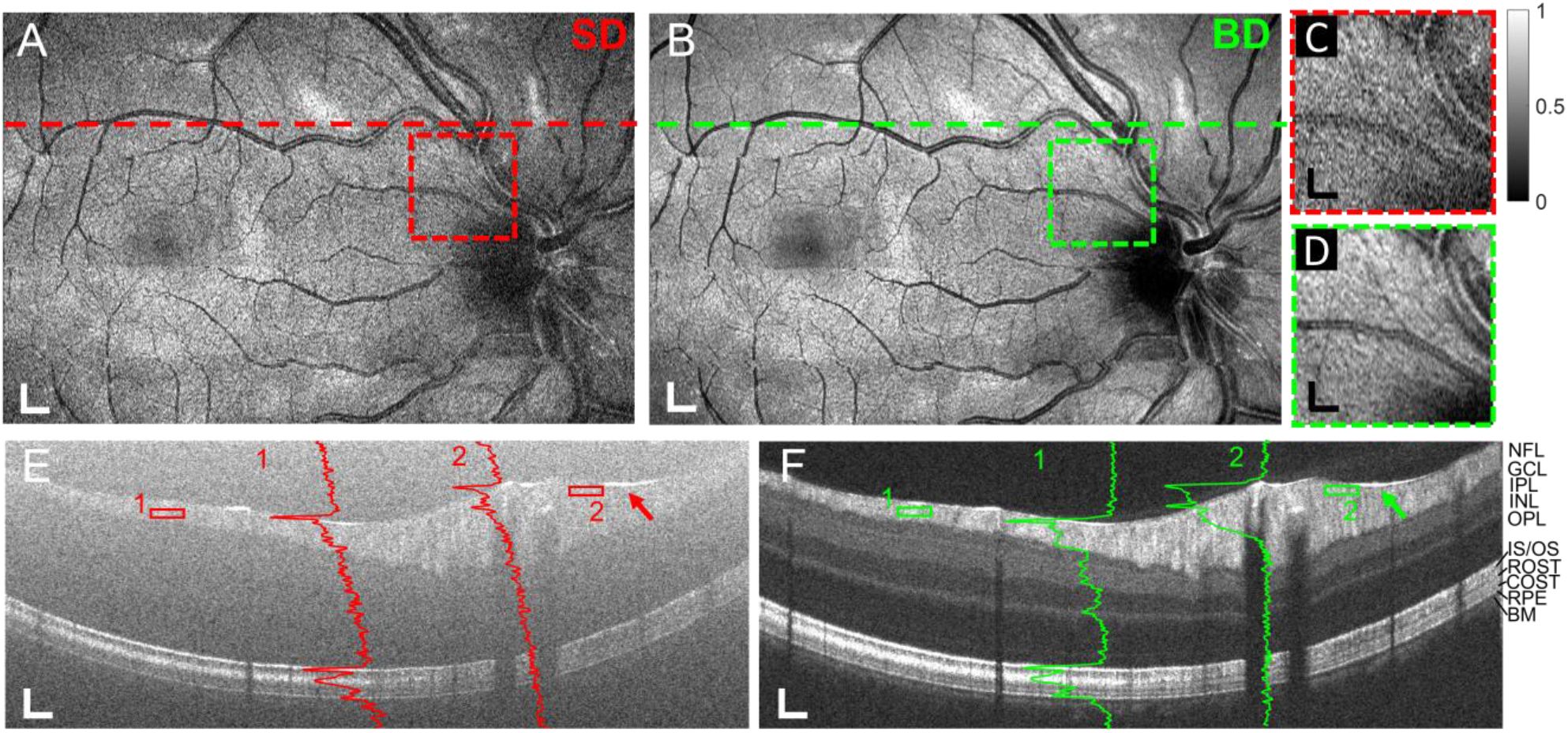
(A-F) Medium field-of-view vis-OCT images of the retina of a healthy 24-year-old male (Eye 2); (A) *En face* projection near fovea for single detection; (B) *En face* projection near fovea for balanced detection; (C) Magnification from red dashed box in (A); (D) Magnification from green dashed box in (B); (E) B-scan from location of dashed line in (A); (F) B-scan from location of dashed line in (B); Red and green A-lines overlay their respective locations; Red and green arrows highlight locations of *SNFR* measurement; Solid red and green boxes highlight locations of *CNR* measurement. Scale bars in (A & B) are 275 *μ*m (vertical) × 325 *μ*m (horizontal); scale bars in (C & D) are 150 *μ*m (vertical) × 175 *μ*m; scale bars in (E & F) are 50 *μ*m (vertical) × 275 *μ*m (horizontal). NFL: nerve fiber layer; GCL: ganglion cell layer; IPL: inner plexiform layer; INL: inner nuclear layer; OPL: outer plexiform layer; ELM: external limiting membrane; IS/OS: inner segment/outer segment; COST: cone outer segment tips; ROST: rod outer segment tips; RPE: retinal pigment epithelium; BM: Bruch’s membrane

**Figure 6.**
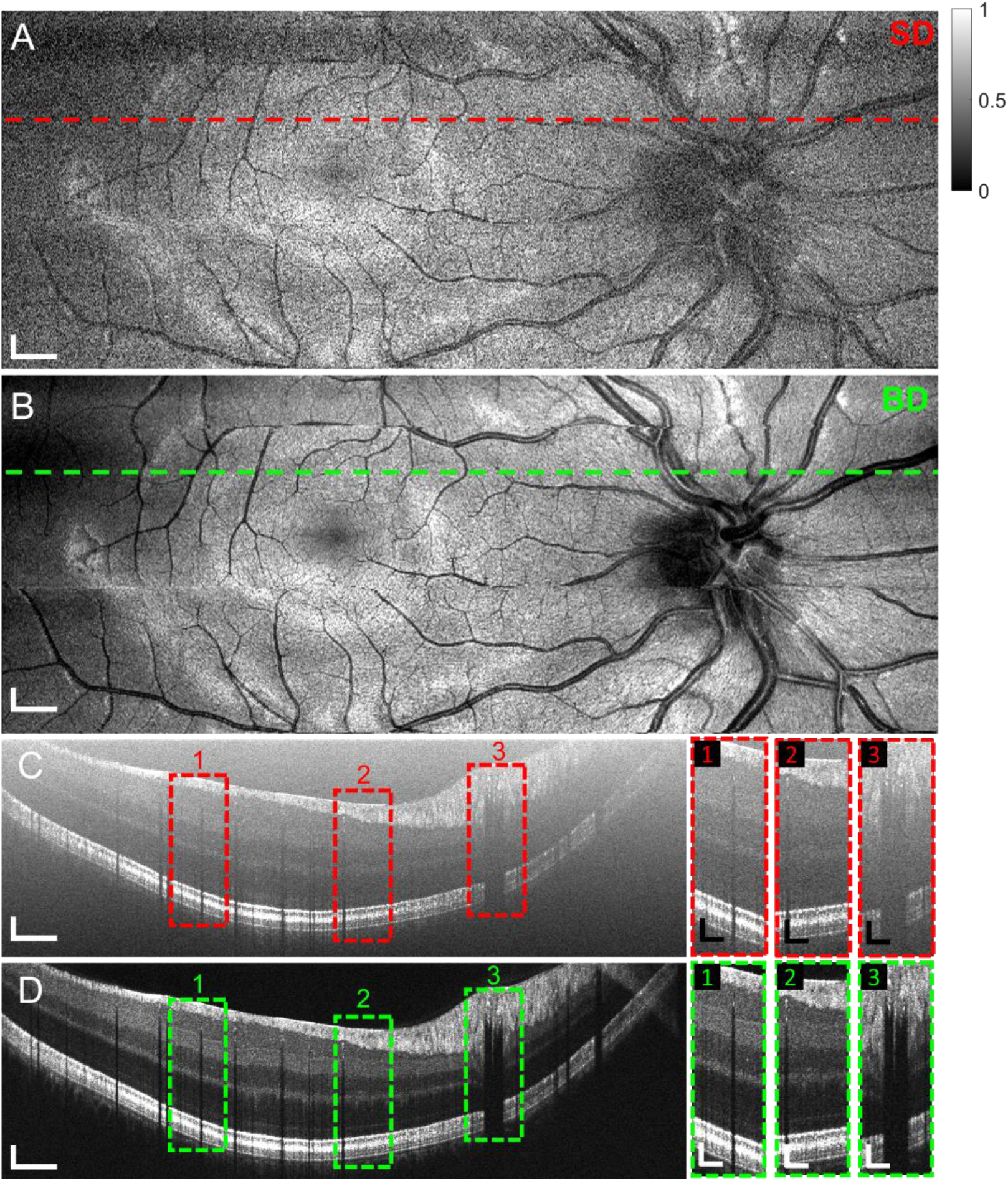
(A-D) Large FOV scans in 24-year-old male (Eye 2); (A) *En face* projection for balanced detection; (B) *En face* projection for single detection; (C) High-density speckle reduction (HDSR) scan at location of red dashed line in (A); (D) High-density speckle reduction (HDSR) scan at location of green dashed line in (B). Red and green dashed boxes in (C) & (D) labeled 1-3 highlight regions for magnification in SD and BD, respectively. Magnifications are shown to right and labeled 1-3 for SD and BD, respectively. Scale bars in (A & B) are 260 *μ*m (vertical)× 500 *μ*m (horizontal); scale bars in (C & D) are 40 μm × 700 μm; scale bars in magnification boxes 1-3 are 40 μm × 200 μm

## Results

### Noise cancelation

We validated BD-vis-OCT by comparing the noises in the spectrometers before and after BD from three locations in the 1D CCD camera array. Fig. 2 shows noises used to calibrate the spectrometers at the pixel numbers 400, 950, and 1550, which represent the starting, middle, and ending positions of the 2048 camera pixels. Fig. 2A plots noises from 100 consecutive acquisitions at pixel 400 detected by SRA (blue) and SRB (orange) after DC removal. The standard deviation (*σ*, [arb. Units]) of noises are 97.6 for SRA and 104.9 for SRB. Despite the BD configuration, SRA and SRB still did not have identical k-sampling (Fig. 1C), and therefore did not measure the same noises. This is confirmed qualitatively by the lack of overlap in the noise measured from SRA (blue line) and SRB (orange line) in Fig. 2A. Quantitatively, the noises have a relatively low correlation coefficient (CC) of 0.47. Fig. 2B plots the same noises after calibration. After calibration, the noises well overlap with a CC of 0.98. Fig. 2C plots the difference between the noises from SRA and SRB with and without calibration. The noise subtraction without proper calibration (plotted in yellow) has a significantly higher standard deviation (*σ* = 104.2) than that after calibration (*σ* = 22.6) (plotted in purple). We validated these trends for each pixel in the spectrometer. Figs. 2D-2F and Figs. 2G-2I plot the same comparisons as Fig. 2A-2C for pixels 950 and 1550, respectively. For pixel 950, SRA had *σ* = 116 and SRB had *σ* = 105.5. The CC was 0.42 without calibration and 0.96 with calibration. Subtraction yielded *σ* = 119.6 without calibration and *σ* = 28.5 with calibration. For pixel 1550, SRA had *σ* = 70.7 and SRB had *σ* = 64.7. The CC was 0.31 without calibration and 0.99 with calibration. Subtraction yielded *σ* = 79.4 without calibration and *σ* = 14.4 with calibration. The average CC across all pixels was 0.96 with calibration and 0.39 without calibration.

### Imaging phantom eyeball

We imaged a phantom eyeball to assess BD-vis-OCT retinal imaging in a well-controlled environment. We compared SD and BD images using the camera’s lowest and highest gains and quantified *SNFR*, *G*, and *CNR*.

Figs. 3A & 3B show SD and BD B-scans (five-times averaging) of the phantom at the lowest camera gain. The images are plotted on the same contrast scales (square root, normalized between 0 and 1). While the SD image failed to resolve the phantom structure clearly, the BD image fully revealed all the structure details. Fig. 3C plots the A-lines from the locations highlighted by the red and green dashed lines in Figs. 3A and 3B, respectively. The peak near 220 *μ*m depicts the bright layer on the top of the phantom. The peak from BD (green line) is 5 dB higher than SD (red line). As explained above, the perceived *G* can be less than 6 dB due to a bias at very low signal-to-noise ratios. The noise floor is located to the left of the peak at depths near 150 *μ*m. At this location, the noise floor is 17 dB lower for BD than SD. The SD image has an average *SNFR* of 5.9 dB, and the BD image has an average *SNFR* of 28.1 dB, resulting in an average *G* of 22.2 dB. We measured *CNR*s in the boxes highlighted by 1-3 in Figs. 3A & 3B, respectively. They are −8.3 dB, −9.8 dB, and noise floor limited for SD, and 4 dB, 3.8 dB, and 4.1 dB for BD, respectively.

Figs. 3D & 3E show SD and BD B-scans (five-times averaging) of the phantom at the highest camera gain. The images are plotted on the same contrast scales (square root, normalized between 0 and 1). With a higher gain, it takes less reference power to saturate the camera, thereby reducing the contribution of RIN to the SD A-line [28]. Fig. 3C plots A-lines from the regions highlighted by the red and green dashed lines in Figs. 3A and 3B, respectively. The peak from BD (green line) is 6 dB higher than that from SD (red line), consistent with the maximum theoretical increase. The noise floor is 9 dB lower for BD than SD. The SD image has an average *SNFR* of 12.4 dB, and the BD image has an average SNFR of 27.3 dB, an average increase of 14.9 dB. We measured CNRs in the boxes labeled 1-3 in Figs. 3A & 3B, respectively. They are 0.1 dB, 0.8 dB, and 1.7 dB for SD, and 3.7 dB, 3.5 dB, and 4 dB for BD, respectively.

For SD, the average *SNFR* is 6.5 dB lower at the lowest camera gain (5.9 dB) than at the highest camera gain (12.4 dB). CNR is noise floor limited at the lowest camera gain and 1.7 dB at the highest camera gain. This difference is attributed to RIN. Nevertheless, for BD, the *SNFR*s (28.1 dB and 27.3 dB) and *CNR*s (4.1 dB and 4 dB) under the lowest and highest camera gains. Therefore, BD generates similar image qualities at both gains, despite the SD image qualities being vastly different.

### Human retina images – small FOV

Figs. 4 shows the small FOV scan located near the fovea of the right eye of a healthy, 47-year-old male volunteer (Eye 1). Figs. 4A & 4B show *en-face* images for SD and BD, respectively. We generated the *en-face* images by taking the mean intensity projection of the retinal volume after cropping out the first 20 pixels along the depth direction. The red and green dashed boxes in Figs. 4A & 4B highlight regions that are magnified by Figs. 4C and 4D, respectively. Fig. 4D highlights small vessels buried in the noise floor without BD, as shown in Fig. 4C. Figs. 4E & 4F show B-scans (registered and averaged five times) at the locations highlighted by the red and green dashed lines in Figs. 4A & 4B, respectively. Comparing with SD (Fig. 4E), where the high noise floor makes the retinal anatomical features hard to be distinguished, the BD B-scan image resolved all the fine anatomical details [10, 29] against the background, as highlighted in Fig. 4F. To further compare the difference in structural visibility between SD and BD, we overlaid A-lines (log scale) on their respective locations in Figs 4E and 4F. A-line 1 highlights a blood vessel at its respective location in the SD (red A-line) and BD (green A-line). For SD, the signal within the blood vessel is buried in the noise floor. Meanwhile, the reduced noise floor in BD reveals the characteristic blood signal decay [30, 31]. The attenuation is visible across the entire depth of the vessel. A-line 2 highlights the retinal anatomical layers resolved by SD and BD, respectively. BD reveals the inner plexiform layer (IPL) sublayers, which has three distinct bright laminations and is a promising biomarker for glaucoma [11, 12]. In SD image and A-line, the IPL is invisible. We measured SNFRs at an ILM reflection (highlighted by red and green arrows in Figs. 4E & 4F, respectively). The SNFR is 10.9 dB in SD and 25.8 dB in BD, resulting in a *G* = 14.9 dB. We measured CNRs in the RNFL highlighted by the red and green boxes labeled 1 and 2 in Figs. 4E & 4F. In SD, CNRs are −2.6 dB and noise floor limited for boxes 1 and 2, respectively. In BD, CNRs are 5.2 dB and 5.5 dB for boxes 1 and 2, respectively.

There are minimal motion artifacts in the small FOV volume. We did not perform any registrations for the *en face* projections. At 125-kHz A-line rate, the small FOV volumes were acquired in 1 second, overcoming many fundamental limitations from eye motions, where, for example, involuntary saccades occur on the order of 1 Hz. Additional images with small FOV from other volunteers are shown in **Supplementary Information**.

### Human retina imaging with medium FOV

Fig. 5 compares SD and BD vis-OCT image with medium FOV from the left eye of a 24-year-old male volunteer (Eye 2). *En face* projections (Figs. 5A and 5B) cover the fovea and optic disc (OD) in a single scan with minimal motions. In the BD image (Fig. 5B), as compared with SD, vessels from the OD to the fovea are visible at the capillary level thanks to the improved optical contrast within the visible-light spectral range. Figs. 5C and 5D are magnified views of the areas highlighted by the red and green dashed boxes in Figs. 5A and 5B, respectively, where capillaries are better visualized by BD.

Figs. 5E and 5F show B-scans (registered and averaged five times) from respective locations highlighted by the red and green dashed in Figs. 5A and 5B. Similar to the small FOV B-scans, BD reveals the retinal anatomical layers with much higher contrast than SD. In previous work by Rubinoff et al. [9], we showed that B-scans acquired at an A-line rate of 25 kHz could not be registered and directly averaged without significant blurring. Here, we show that with a 125 kHz A-line rate, five B-scans can be registered and averaged with nearly no blurring. We previously limited vis-OCT’s A-line rate at 25 kHz to reduce the influence of RIN and increase image quality in human imaging. BD achieved comparable image quality at a 5-fold increased A-line rate. In Figs. 5E and 5F, we overlay two A-lines (log scale) that highlight retinal structures and a blood vessel at their respective locations (highlighted 1 & 2). We measured SNFR at the regions highlighted by the red and green arrows, respectively. We found that SNFRs were 16.2 dB and 31.4 dB for SD and BD, respectively, resulting in a *G* = 15.2 dB. We measured CNR in the areas highlighted by the red and green boxed regions labeled 1 and 2, respectively. In Fig. 5E, CNRs are −1.6 dB and noise floor limited for boxes 1 and 2, respectively. In Fig. 5F, CNRs are 5 dB and 4.7 dB for boxes 1 and 2, respectively. Additional images with medium FOV from other volunteers are shown in **Supplementary Information**.

### Human retinal imaging with large FOV and high density speckle reduction

Figs. 6A and 6B respectively show *en-face* SD and BD vis-OCT images from Eye 2 with large FOV and Figs. 6C and 6D respectively show high-density speckle reduction (HDSR) [9] SD and BD B-scan image from the positions highlighted by red and green dashed lines in Figs 6A and 6B, respectively. The large FOV scan enables vis-OCT to provide an unprecedented view of the retina beyond the macula and optic disc while exhibiting minimal motions thanks to the 125-kHz A-line rate. We combined the large FOV scan with an HDSR scan. A benefit of the HDSR scan is that it can combine dense A-line scanning, speckle reduction, and wide scanning range without registrations [9]. Dense A-line scanning can help reduce fringe washout from the scanner. At the 125 kHz A-line rate with BD, we can increase HDSR A-line density without sacrificing the total scanning time. Here, we acquired HDSR B-scans with 32768 A-lines and laterally averaged them 32 times, allowing many more averages than in the medium FOV (five averages). The red and green dashed boxes (1-3) in Figs. 6C and 6D highlight three retinal regions in the HDSR B-scan image, and their magnified views are shown on the right. Following the pattern in Figs. 3–5, BD, but not SD, reveals the structure (boxes 1 & 2) and blood vessels (box 3) across the retina. As seen in Fig. 6D and magnified by boxes 1 & 2, IPL, RPE, and BM (see Figs. 4 & 5) are delineated across nearly the entire scan range. vis-OCT imaging across a large FOV may be useful for mapping structures like IPL, RPE, and BM, leading to improved diagnosis, monitoring, and treatment of blindness causing diseases like glaucoma and macular degeneration. An additional example of an HDSR scan is available in **Supplementary Information**.

## Discussion

We demonstrated BD in vis-OCT for the first time after a precise calibration of two spectrometers using the RIN itself. We showed in a phantom eyeball that BD reaches similar SNFRs under both maximal (27.3 dB) and minimal (28.1 dB) camera gains, despite SD SNFR being significantly lower under the minimal camera gain. This implies not only that BD is highly effective in removing RIN even at low SNFRs, suggesting vis-OCT can afford shorter spectrometer exposure times and higher A-line rates. In the future, we plan to conduct systemic investigations of the influence of spectrometer detection gain on BD performance. Based on our *in vivo* results, we anticipate that BD-vis-OCT can reach A-line rates higher than 200 kHz, although one potential limitation is fringe washout caused by optical scanning.

We found *in vivo* in four volunteers aged 25-47 that *G* is near the theoretical limit found in the retinal phantom (~ 15 dB). We found *CNR* increases from noise floor limited to up to 5.5 dB. Most importantly, we found that retinal features entirely obscured by RIN in SD are revealed by BD. Based on the image comparisons, we did not observe any significant artifacts or resolution loss that may be attributed to poor calibration.

We noted that SD *en-face* projections, although noisy, appear less obscured by noise than the corresponding B-scans. This may seem counterintuitive at first since most of the contrast in the *en face* projections originate from large vessels located near the inner retina, which are mostly obscured by the noise floor in their corresponding B-scans. This is because significant vessel contrast in the *en face* projections is derived from shadows created by blood attenuation, not the blood signal itself. In vis-OCT, blood’s attenuation coefficient is more than 10-fold higher than in NIR OCT [32]. To that end, the vessel shadows create high contrast against the brightly reflecting photoreceptors and RPE, which may be observable even in SD. We show this in **Supplementary Information** (Fig. S4), where the vessels in the SD *en face* projection (Fig. S4A) are mostly obscured by the noise when photoreceptor layers in the B-scan (Fig. S4E) are not clearly visible. It is notable that three-dimensional (3D) vessel maps of the retina, which may be used for oximetry, cannot rely on the shadow effect and must have sufficient vessel signal.

RIN was previously suppressed in vis-OCT by reducing the A-line rate to perform increased temporal averaging of the interference fringe. When combined with the distraction and discomfort of a visible-light beam, low speed is perhaps the critical limiting factor for translating vis-OCT to the clinics. BD enables high-speed vis-OCT without sacrificing image quality, resolution, or spectroscopic bandwidth. Additionally, supercontinuum lasers are significantly more costly than SLDs, and technology improvements to reduce RIN in supercontinuum lasers are likely to add more costs. BD-vis-OCT eliminates the influence of RIN, permits high-speed, high-quality human imaging, and imposes fewer performance requirements on supercontinuum lasers, all of which will likely bring down the overall vis-OCT cost. Finally, although we focus on vis-OCT here, it is not the only OCT technology where BD is applicable. Other OCTs using large bandwidths to increase axial resolution [33], or those operating in the far-infrared range to achieve deep light penetration [34], also rely on supercontinuum sources.

## Conclusion

We developed BD-vis-OCT based on an MZI. Using BD, we demonstrated high-quality human retinal imaging at an A-line rate of 125 kHz. We demonstrated sensitivity gains up to 22.2 dB in a phantom and up to 15.2 dB in human eyes. The majority of *SNFR* gains came from the reduction of RIN. Qualitatively, BD revealed more structural features *in vivo* than SD. We anticipate that BD will enable broader clinical applications of vis-OCT.

## Acknowledgments

The authors thank Lisa Beckmann for her helpful discussions. The authors are grateful for the National Institutes of Health grants R01EY019949, R01EY026078, R01EY028304, R01EY029121, and R44EY026466.

## Supplementary Information

### Axial resolution measurement for single and balanced detections

**Figure S1.**
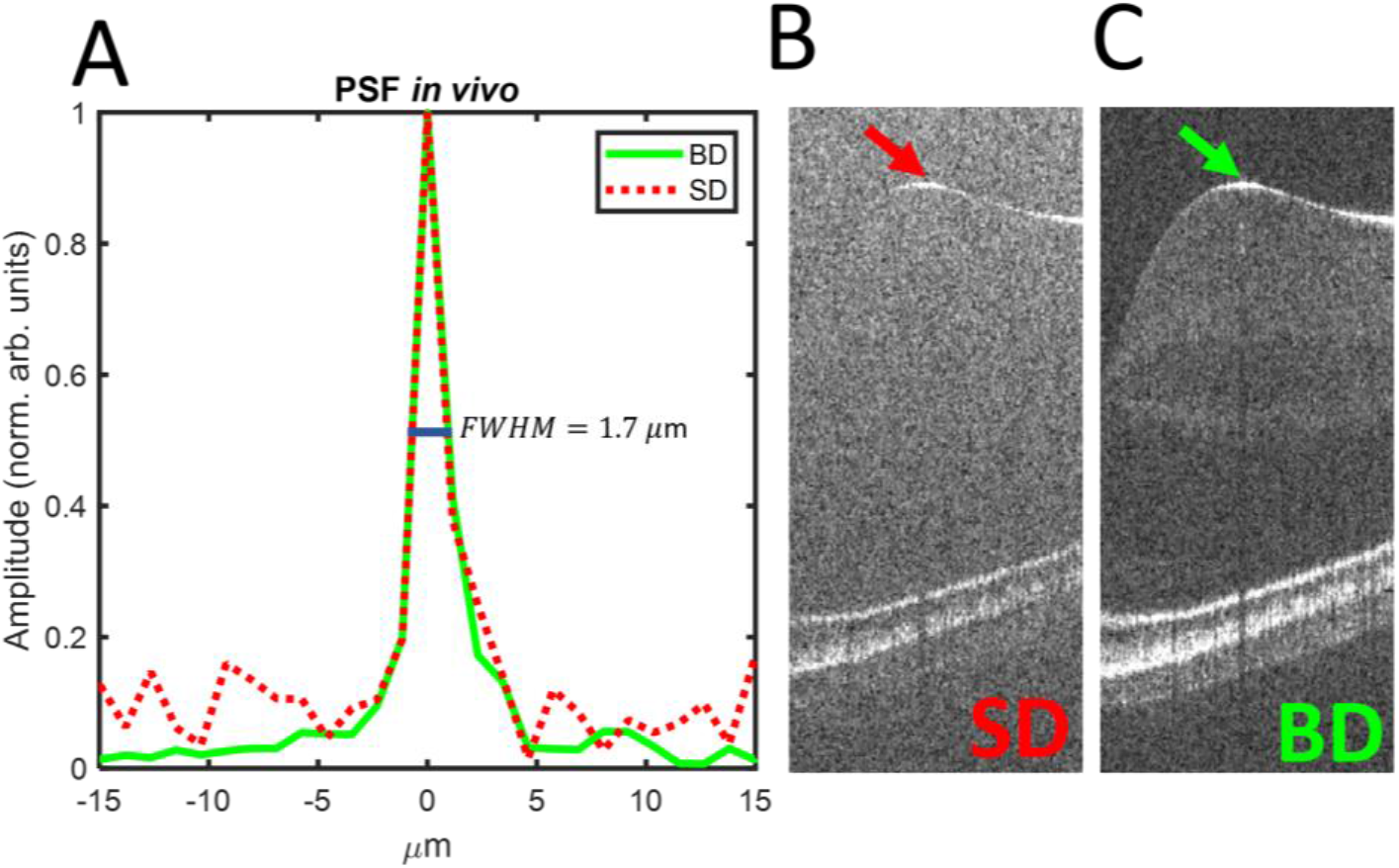
Resolution measurement for BD and SD; (A) Point spread functions (PSF) from specular reflection at ILM. Solid green line represents PSF from BD and red dashed line represents PSF from SD. Both PSFs overlap and have a measured full-width and half maximum (FWHM) of 1.7 *μ*m. (B) *En face* projection single detection; (B) Location of PSF for SD (highlighted by red arrow); (C) Location of PSF for BD (highlighted by green arrow)

### Additional examples of *in vivo* retinal imaging

**Figure S2.**
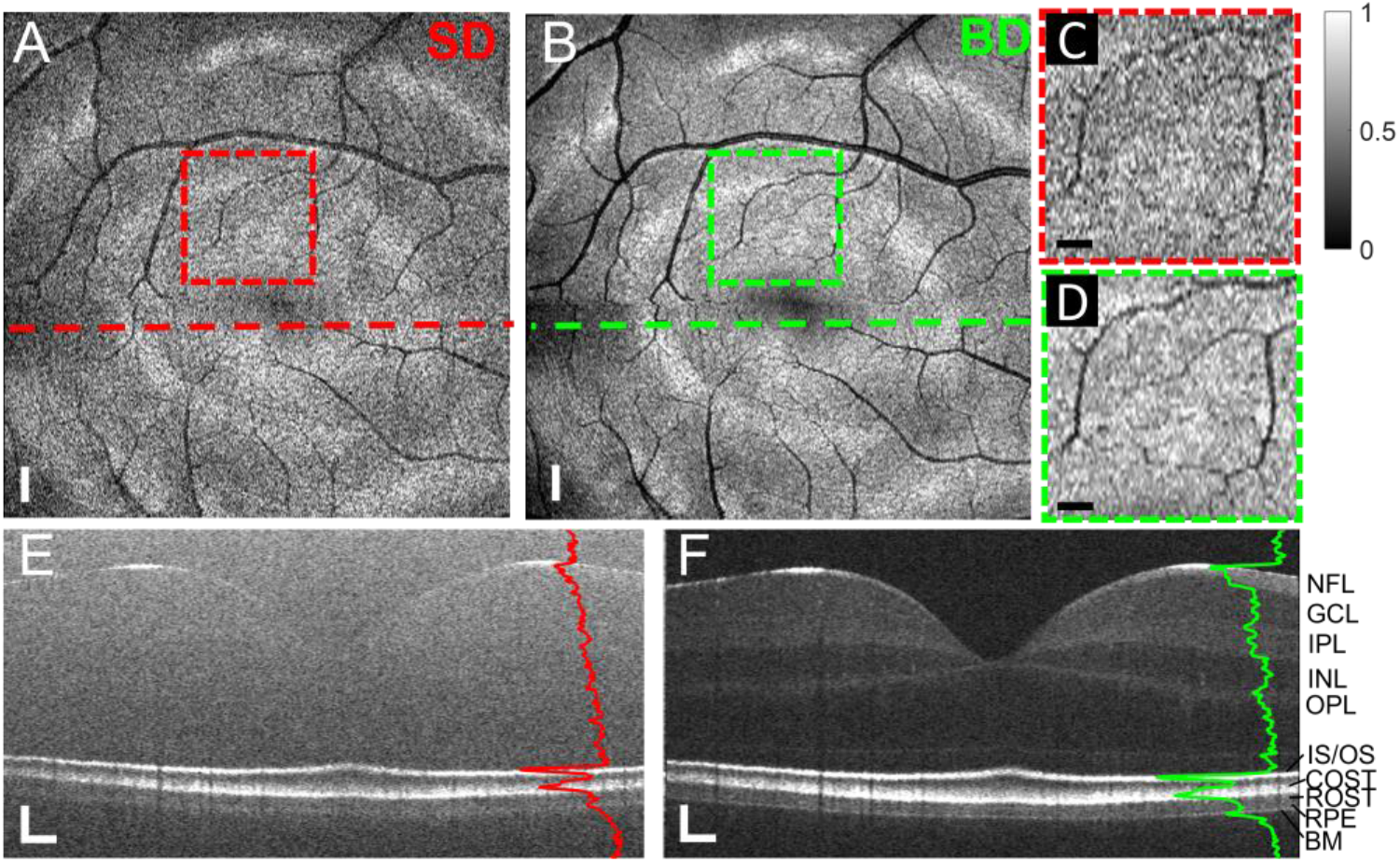
(A-F) Small field-of-view vis-OCT images of the retina of a healthy 24-year-old male (Eye 2); (A) *En face* projection near fovea for single detection; (B) *En face* projection near fovea for balanced detection; (C) Magnification from red dashed box in (A); (D) Magnification from green dashed box in (B); (E) B-scan from location of dashed line in (A); (F) B-scan from location of dashed line in (B); Red and green A-lines overlay their respective locations Scale bars in (A & B) are 275 *μ*m (isometric); scale bars in (C & D) are 150 *μ*m (isometric); scale bars in (E & F) are 60 *μ*m (vertical) × 225 *μ*m (horizontal). NFL: nerve fiber layer; GCL: ganglion cell layer; IPL: inner plexiform layer; INL: inner nuclear layer; OPL: outer plexiform layer; ELM: external limiting membrane; IS/OS: inner segment/outer segment; COST: cone outer segment tips; ROST: rod outer segment tips; RPE: retinal pigment epithelium; BM: Bruch’s membrane

**Figure S3.**
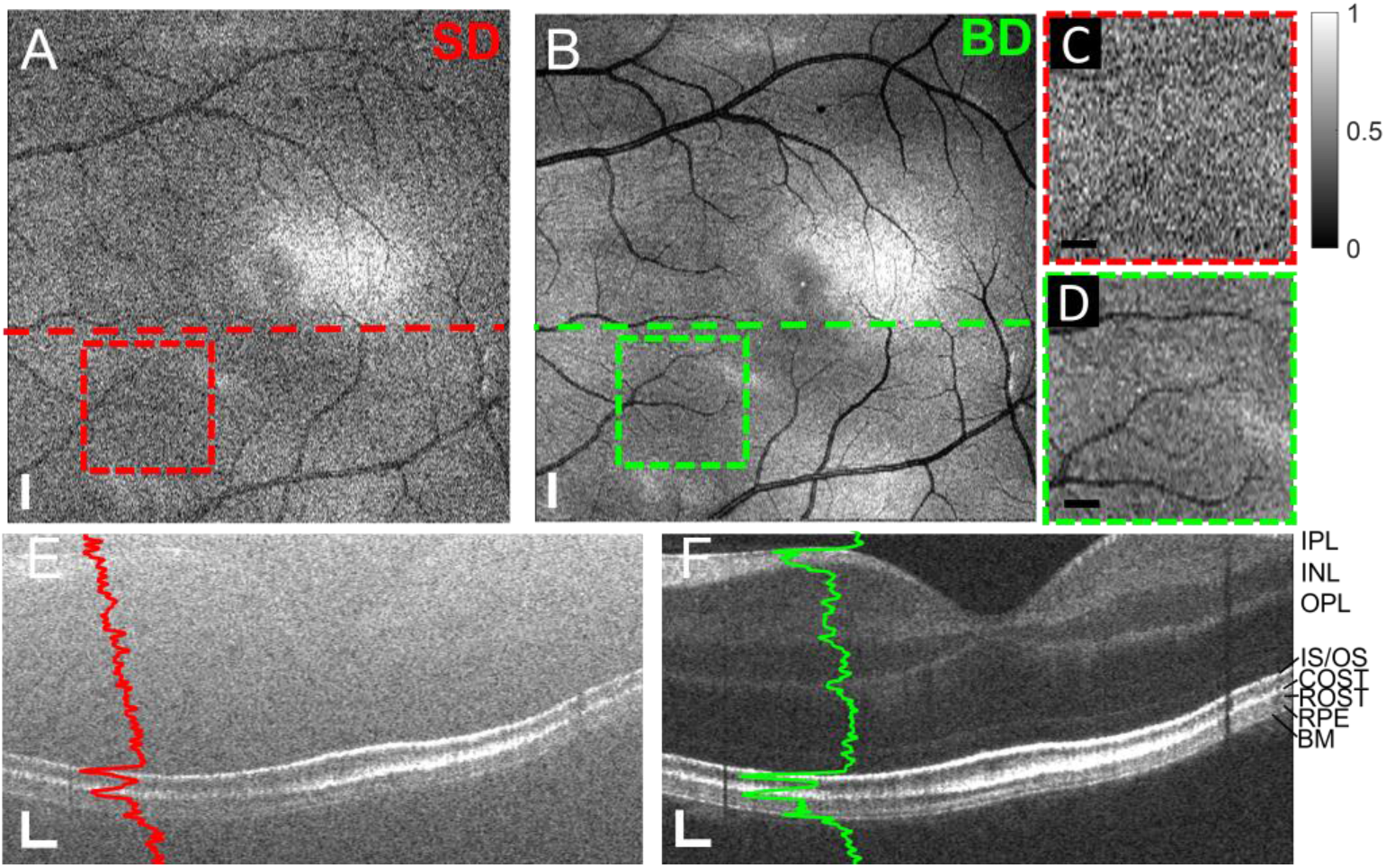
(A-F) Small field-of-view vis-OCT images of the retina of a healthy 27-year-old female (Eye 3); (A) *En face* projection near fovea for single detection; (B) *En face* projection near fovea for balanced detection; (C) Magnification from red dashed box in (A); (D) Magnification from green dashed box in (B); (E) B-scan from location of dashed line in (A); (F) B-scan from location of dashed line in (B); Red and green A-lines overlay their respective locations. Scale bars in (A & B) are 275 *μ*m (isometric); scale bars in (C & D) are 150 *μ*m (isometric); scale bars in (E & F) are 60 *μ*m (vertical) × 225 *μ*m (horizontalIPL: inner plexiform layer; INL: inner nuclear layer; OPL: outer plexiform layer; ELM: external limiting membrane; IS/OS: inner segment/outer segment; COST: cone outer segment tips; ROST: rod outer segment tips; RPE: retinal pigment epithelium; BM: Bruch’s membrane

**Figure S4.**
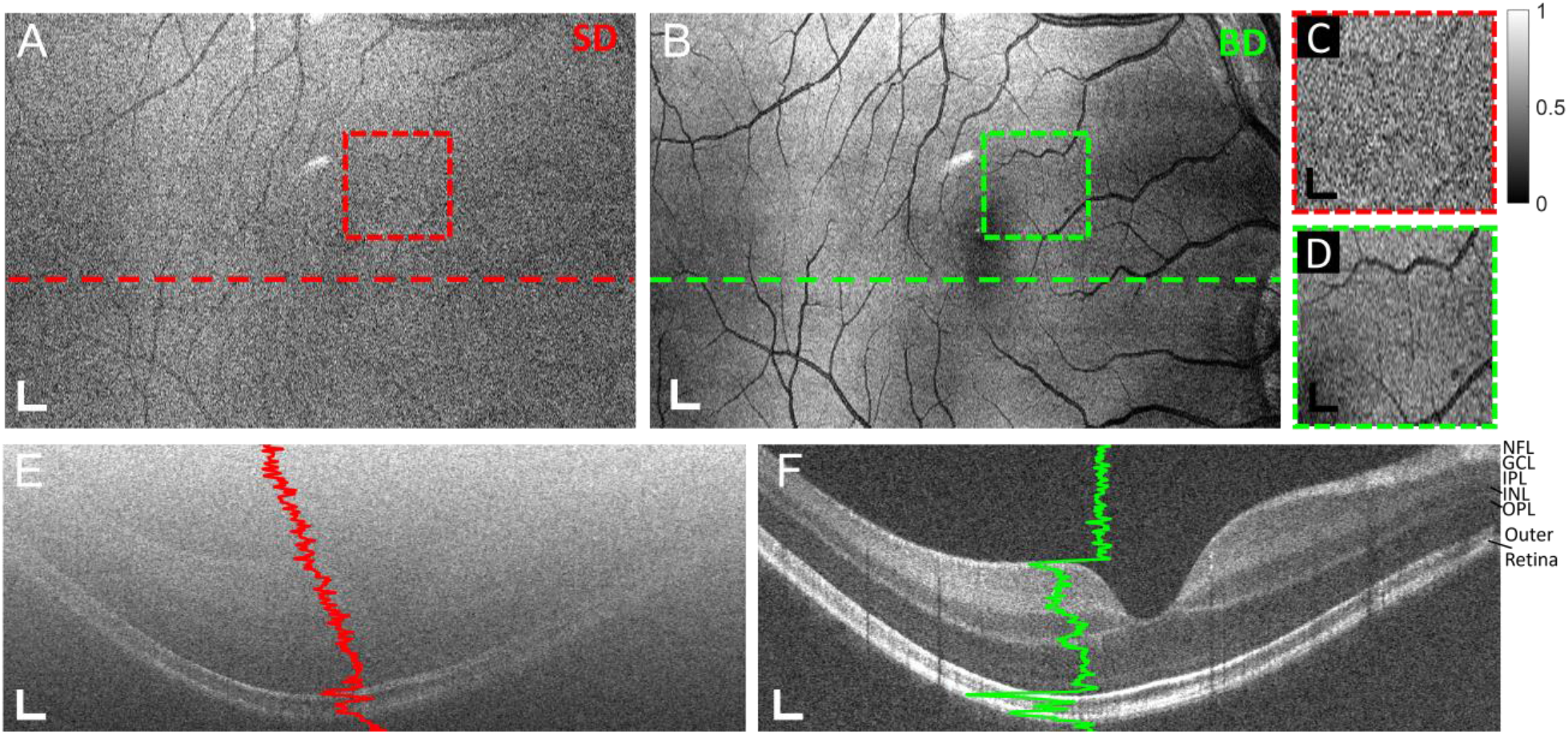
(A-F) Medium field-of-view vis-OCT images of the retina of a healthy 25-year-old male (Eye 4); (A) *En face* projection near fovea for single detection; (B) *En face* projection near fovea for balanced detection; (C) Magnification from red dashed box in (A); (D) Magnification from green dashed box in (B); (E) B-scan from location of dashed line in (A); (F) B-scan from location of dashed line in (B); Scale bars in (A & B) are 365 *μ*m (horizontal) × 275 *μ*m (vertical); scale bars in (C & D) are 200 *μ*m ×150 *μ*m; scale bars in (E&F) are 50 *μ*m × 275 *μ*m. B-scans magnified to show detail. NFL: nerve fiber layer; GCL: ganglion cell layer; IPL: inner plexiform layer; INL: inner nuclear layer; OPL: outer plexiform layer

**Figure S5.**
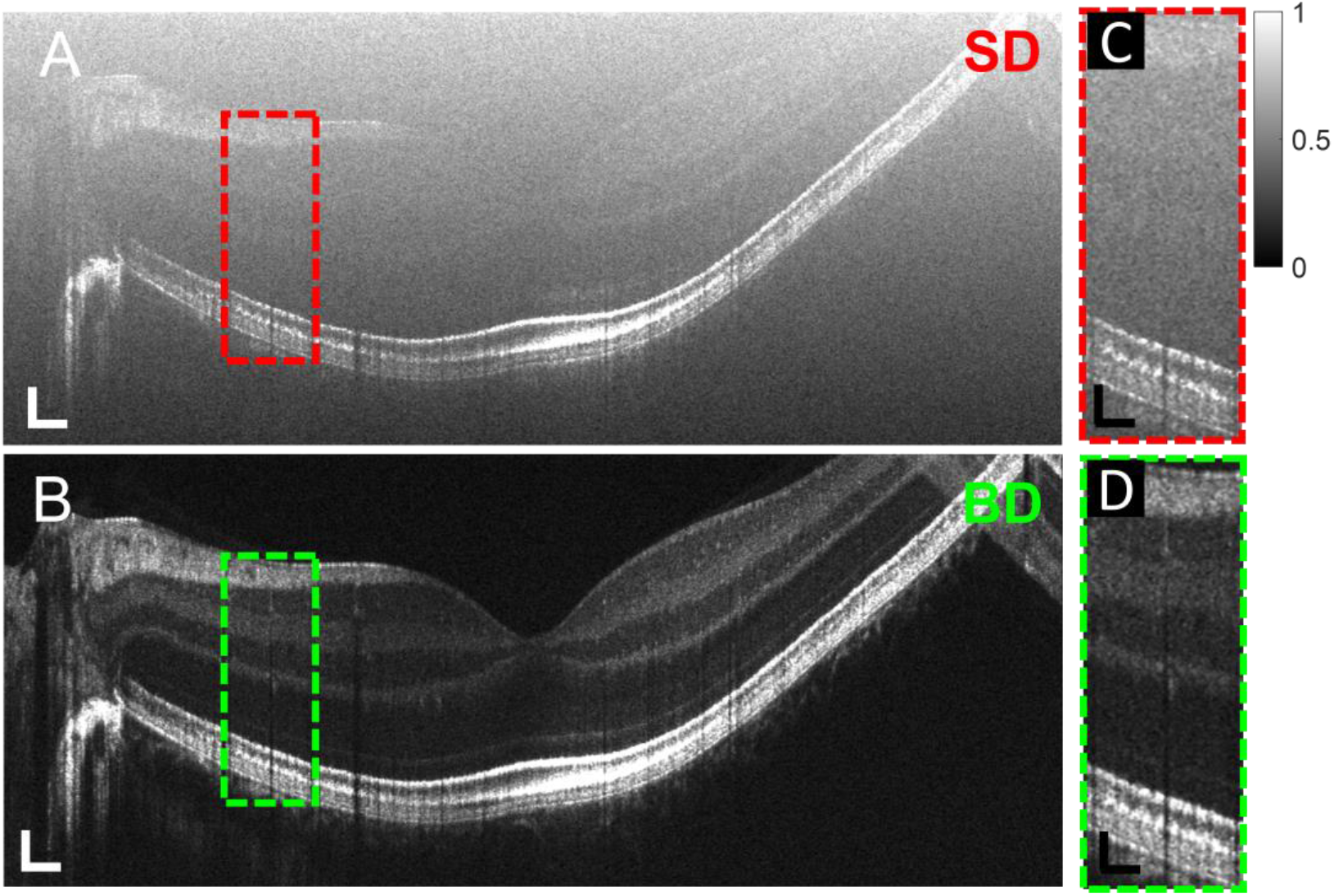
(A-D) High-definition speckle reduction (HDSR) vis-OCT image of the retina of a healthy 27-year-old female (Eye 3). (A) B-scan for SD; (B) B-scan for BD; (C) Magnification from red dashed box in (A); (D) Magnification from green dashed box in (B); Scale bars in (A & B) are 60 *μ*m (vertical)× 235 *μ*m (horizontal); scale bars in (C & D) are 70 μm × 150 μm;

